# Age-dependent Powassan Virus Lethality and Neuropathogenesis in Mice

**DOI:** 10.1101/2023.05.11.540417

**Authors:** Megan C. Mladinich, Grace E. Himmler, Jonas N. Conde, Elena E. Gorbunova, William R. Schutt, Stella Tsirka, Hwan Keun Kim, Erich R. Mackow

**Affiliations:** Department of Microbiology and Immunology, Center for Infectious Disease, Stony Brook, New York, USA; Department of Pharmacological Sciences Stony Brook University, Stony Brook, New York, USA

**Author notes:** Corresponding Author Erich R. Mackow, Ph.D., AAM and AAAS Fellow, Department of Microbiology and Immunology, Renaissance School of Medicine, Life Sciences Rm 126, Stony Brook University, Stony Brook, NY 11794-5222.

## Abstract

Powassan viruses (POWV) are emergent tick-borne flaviviruses that cause severe neurologic disease in humans. Subcutaneous inoculation of C57BL/6 mice with POWV (strain LI9) resulted in overt brain damage resembling spongiform encephalitis. Noting higher POWV lethality in older mice, we assessed neurovirulence as a function of age. We found that POWV LI9 inoculation was lethal in 80% of 50 wk old mice, 10-15 dpi, and that lethality was sequentially reduced in 40, 30, 20, 10 wk old mice to <10%. Lethality was conferred by 2-20 POWV FFUs, and POWV neuropathology was evident as early as 5 dpi, with lethal disease 10-15 dpi correlated with sustained POWV RNA levels in brains of aged mice. Histology of POWV infected 50 wk old murine brains revealed severe spongiform neuronal necrosis, microgliosis, and inflammation with increased brainstem and cerebellar damage. These findings delineate an age-dependent murine model of lethal POWV infection that mirrors human POWV disease and permits analysis of age-dependent neurovirulence determinants.

**Significance:** Our findings establish a novel age-dependent lethal animal model to study encephalitic POWV disease *in vivo*. These initial findings demonstrate that following peripheral inoculation, non-neuroadapted POWV LI9 is neuroinvasive and enters the brains of young and aged mice. However, POWV LI9 lethality is strictly age-dependent and correlated with increased viral load in the brains of aged mice. POWV rapidly directs neuronal loss and spongiform lesions, microglial activation and causes prolonged inflammation that fails to clear POWV from the brains of aged mice. Our results provide a lethal murine model of POWV neurovirulence that mirrors the prevalence of severe human POWV encephalitis in the elderly. This lethal murine POWV model provides mechanisms for defining POWV protective responses of the young, revealing determinants of age-dependent POWV lethality and evaluating potential POWV therapeutics.

**SUMMARY:** Powassan virus is an emerging tick-borne flavivirus linked to severe neurologic disease in aged individuals. Here we describe an age-dependent mouse model of POWV pathogenesis.

**SUBJECTS:** Powassan virus, flavivirus, neurovirulence, neuroinvasion, neurotropic, spongiform encephalopathy, microgliosis, neuroinflammation

## INTRODUCTION

Flaviviruses (FVs) are a family of positive-strand RNA viruses that cause wide-spread human disease. Tick-borne FVs (TBFVs) cause up to 15,000 annual cases of severe encephalitis^[1]^, and Powassan virus (POWV) is an emerging neurotropic TBFV primarily spread by deer ticks (*I. scapularis*) in the Northeastern United States^[2, 3]^. POWV is rapidly transmitted in tick saliva during a 15-minute bite, and the prevalence of POWV infections has increased due to improved surveillance and the geographic expansion of tick vectors^[1, 2, 4-6]^. The lack of POWV vaccines and therapeutics make POWV a growing public health concern.

POWV causes severe encephalitis with a 10% case fatality rate, and long-term neurological morbidities in 50% of survivors^[1]^. Neurotropic FVs, including POWV, Tick-borne encephalitis virus (TBEV), West Nile virus (WNV), Japanese encephalitis virus (JEV), and Zika virus (ZIKV) are characterized by a biphasic disease process^[7-9]^. Cutaneous viral inoculation and hematogenous dissemination result in an acute febrile illness, that is followed by a neuroinvasive stage^[10]^. How neurotropic FVs cross restrictive blood-brain barriers (BBBs) and enters the CNS remains enigmatic, but the severity of encephalitic disease is dependent on both FV infection of neuronal cells and inflammatory CNS responses^[7, 11, 12]^. Neuroinvasion is characterized by increased inflammatory responses (e.g. TNFα, IL-1β, IFNγ, CCL2), the activation of resident CNS microglia^[9, 12-14]^ and cytotoxic CD8+ T infiltrates in brain parenchyma in fatal human encephalitis cases^[15-17]^. In humans, the incidence of POWV, WNV and TBEV infections are uniform across age groups, however there is an increased frequency of severe lethal neurologic disease in patients >50 yr old^[1, 2, 18-20]^.

There are 2 POWV genotypes (Lin I and II) that reflect their primary animal and tick hosts, however POWVs are serologically indistinguishable and comprise a single serotype, with 96% amino acid identity in virion Envelope proteins^[21]^. In a 2020 survey, 2% of Long Island, NY *I. scapularis* ticks were found to harbor POWV. POWV Long Island 9 (LI9), a Lin II POWV strain, was isolated from tick homogenates directly into VeroE6 cells and genetically reflects other northeastern POWVs^[22-24]^. POWV LI9 nonlytically infects VeroE6 cells, primary human brain microvascular endothelial cells (hBMEC) and human pericytes. POWV LI9 was found to spread cell-to-cell in epithelial cells forming large infected cell foci in the presence of neutralizing antibodies. In blood-brain barrier (BBB) models, LI9 is released basolaterally from polarized hBMECs and infects abluminal pericytes ^[22]^. Basolateral spread of POWV in hBMECs suggests a mechanism for POWVs to transverse the BBB and enter neuronal compartments, however, the *in vivo* mechanism of POWV entry into the CNS remains to be resolved.

Animal models are critical for understanding FV neurovirulence and resolving mechanisms of neuropathogenesis to prevent lethal patient outcomes. Current POWV animal models use strains that were neuroadapted to replicate in murine brains (LB, SP, IPS1)^[5, 25-28]^. Peripheral infection of immunocompetent mice with POWV LB results in febrile illness 5 to 6 dpi and rapid lethality in >60% of mice (6-9 dpi)^[26, 28-31]^. Variable rates of lethality in 5-14 wk old mice are suggested to reflect murine strains, viral stocks or inoculation site differences^[27]^. In mice inoculated with POWVs SP or IPS1 the onset of illness is delayed (9-10 dpi) with varied mortality rates indicated to be independent of viral dose. The use of murine brain neuroadapted POWVs constrains our understanding of POWV spread from cutaneous inoculation sites and has the potential to bias neurovirulent responses^[5, 21]^.

The emergence of POWVs in highly populated Northeastern States prompted us to isolate currently circulating POWV LI9 from ticks, without neuroadaptation, and to evaluate POWV spread and neurovirulence in mice. We reported that C57BL/6 mice subcutaneously inoculated with POWV LI9 seroconvert and produce cross-reactive POWV neutralizing antibodies^[22]^. Initial studies revealed that subcutaneous LI9 infection of 10-wk-old C57BL/6 mice caused weight loss and clinical neurologic symptoms 7-10 dpi, with 100% of 10-wk-old mice (N=4) recovering 15 dpi. In contrast, 40-wk-old mice continued to lose body weight 11-15 dpi, with 50% of infected mice succumbing to LI9 infection. Increased lethality in 40 versus 10 wk old mice suggested that LI9 lethality may be age-dependent.

Here we evaluated POWV LI9 neurovirulence as a function of age in 10 to 50-wk old C57BL/6 mice. We found that aged C57BL/6 mice s.c. inoculated with POWV LI9 develop lethal neurovirulent disease with overt murine brain damage. LI9 was fatal in 80% of 50 wk old mice 10-15 dpi, and LI9 lethality was reduced in younger mice: 50% lethal in 40 wk old mice, and 10-20% lethal in 10-30 wk old mice. Mice in every age group were infected, seroconverted and elicited similar neutralizing antibody titers. POWV RNA was also detected in brains of all mice, but only the brains of 40 and 50 wk old mice had sustained high POWV RNA loads 10-15 dpi. Consistent with high CNS loads 15 dpi, inflammatory chemokines (IL-1β, CCL2, CCL5, CXCL10) were highly induced in the brains of 50 wk old LI9 infected mice. Collectively, this data suggests that POWV LI9 is neuroinvasive in mice of all ages, and that lethal neurovirulence in 40-50 wk old mice is a function of age and sustained CNS viral load.

Histology and immunohistochemistry analysis of 50 wk old mouse brains s.c. inoculated with POWV LI9 revealed severe neuropathology from 5-15 dpi with notable spongiform encephalopathy, microgliosis, neuronal necrosis, fibrosis and perivascular cuffing. LI9 infected 50 wk old mice brains revealed nissl bodies in neurons 5 dpi, and a dramatic loss of neurons by 5 dpi. Consistent with brain damage, spongiform lesions were present throughout 50 wk old LI9 infected brains with the most pronounced damage in the cerebellum, hindbrain and brainstem. Iba1 staining revealed the recruitment of activated microglia 5-15 dpi and severe microgliosis 15 dpi. These findings suggest that early neuronal damage, prolonged CNS inflammation and viral load are associated with lethality. Our results reveal an age-dependent murine model of lethal POWV neurovirulence that reflects the prevalence of severe lethal disease in elderly patients. This POWV disease model permits us to define responses that prevent lethality in young mice, and age-dependent mechanisms of lethal POWV neurovirulence.

## RESULTS

### POWV LI9 Causes Neurologic Disease in C57BL/6 Mice

POWV LI9 was isolated directly in VeroE6 cells, instead of murine brains, to avoid neuroadaptive changes that may bias spread from peripheral inoculation sites and alter CNS cell targets and responses^[22]^. To assess LI9 spread and neurovirulence in mice we initially inoculated 10 and 40 week old C57BL/6 mice (N=4) peripherally with LI9 (Supp. Fig. 1A-B). Mice were subcutaneously (s.c.) footpad inoculated with 10^3^ PFU of LI9 and assessed for weight loss and neurologic symptoms 1-20 dpi. Both LI9 infected 10 and 40 wk mice lost body weight 7-10 dpi (10-15%), with signs of clinical illness including ruffled fur, hunched posture, and reduced activity (Supp. Fig. 1A). From 11-15 dpi, 10 wk mice recovered initial weight loss and activity without mortality (Supp. Fig. 1A-1B). In contrast, body weights of LI9 infected 40 wk old mice were reduced by 20% 10-15 dpi, with 50% of mice succumbing to LI9 infection 15 dpi (Supp. Fig. 1A-1B). These findings demonstrate that POWV LI9 causes neurologic disease in young and old mice. In contrast, LI9 lethality was restricted to aged mice, similar to the prevalence of POWV encephalitis and mortality in aged individuals.

### POWV LI9 Lethality is Age-Dependent in C57BL/6 Mice

To investigate POWV LI9 age-dependent lethality, we peripherally inoculated 10, 20, 30, 40 and 50 week old C57BL/6 mice (s.c.) with 2 × 10^3^ FFUs of LI9 and assessed lethal neurovirulent disease 1-20 dpi (Fig. 1). In all age groups LI9 infection resulted in weight loss beginning 8 dpi with signs of clinical illness 8-15 dpi. Initial 5-10% weight loss of LI9 infected 10 wk old mice (8-10 dpi) was recovered by 15 dpi (Fig. 1A). However, in LI9 infected 20-50 wk old mice initial weight loss was maintained and increased from 10-15 dpi. Severe 20-25% weight loss was observed in 50-wk-old mice, with 10-15% weight loss in 20, 30 and 40 wk old mice (Fig 1A, N=10-20 per age group). POWV LI9 infected mice developed neurologic symptoms including hindlimb weakness, ataxia, and flaccid paralysis before succumbing to infection 9-15 dpi (Fig. 1B). POWV LI9 was 10% lethal in 10 wk, 20% lethal in 20 wk, 30% lethal in 30 wk and 40% lethal in 40 wk old mice, respectively. The highest mortality was observed in aged, 50 wk old, mice with an 80% mortality rate. We found no significant difference in the morbidity or mortality of male versus female mice. Sera from LI9 infected mice (30 dpi) revealed seroconversion in all age groups with similar POWV neutralizing antibody titers (IC50) in surviving 10-50 week old mice (Fig. 1C). Our findings reveal a sequential increase in the lethality of 10-50 wk old mice inoculated with POWV LI9, and demonstrate that LI9 lethality is age-dependent. This murine model reflects the prevalence severe human POWV disease in elderly patients and provides a means for defining age-dependent determinants of lethal POWV disease.

**Figure 1.**
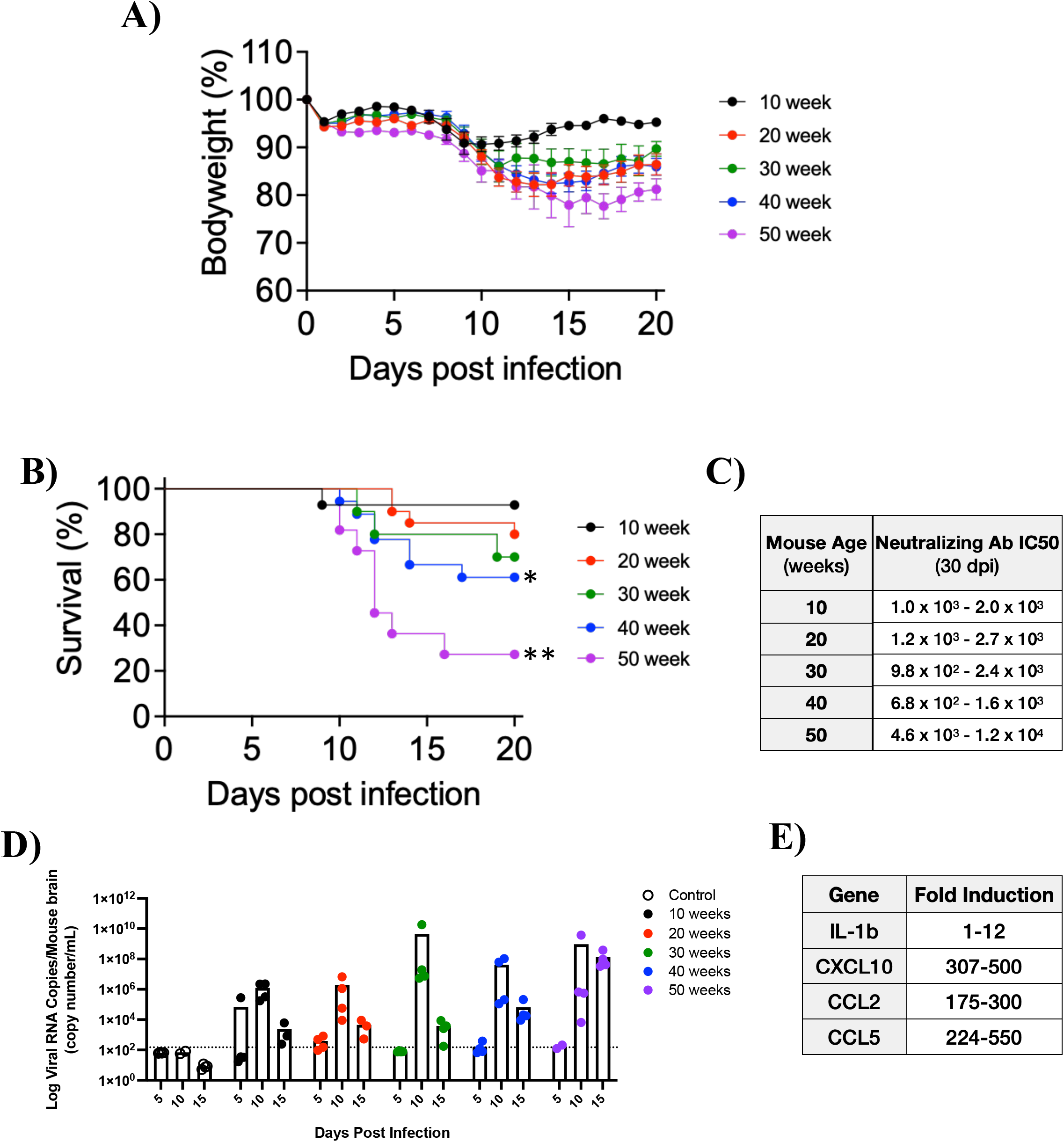
Susceptibility of C57BL/6J Mice to POWV LI9 Increases with Age. (A-E) 10-, 20-, 30-, 40- and 50 wk old mice were inoculated with 2x10^3^ FFU of POWV LI9 or mock-infected with PBS by s.c. footpad injection. (A) Mice were weighed daily, and weights were expressed as a percentage of body weight prior to infection. Results are the mean +/- standard error of the mean (SEM) of the indicated number of mice per group and were analyzed by use of a log rank test. **, *P <* 0.01;*, *P <* 0.05. (B) Lethality was monitored for 20 days. For (A-B): n = 14 (10wk), n = 20 (20wk), n = 10 (30wk), n = 18 (40wk), n = 11 (50wk). (C) Neutralizing antibodies present in POWV LI9 or mock-infected mouse serum were determined by serial dilution and addition to ∼500 FFU of POWV LI9 prior to adsorption to VeroE6 cells. Inoculated VeroE6 cells were washed, and 1 dpi and infected cells were quantitated and IC50 ranges values were determined. For C: n = 4 (10, 20, 30, 40 wk: 30 dpi), n = 3 (50 wk 30 dpi), n = 4 (control). (D) POWV RNA in whole brain was measured at 5, 10, 15 and 30 dpi by qRT-PCR. Data are expressed as copy number milliliter after normalization to a standard curve generated in parallel. For D: n = 4 (10, 20, 30, 40, 50 wk: 5, 10, 15 dpi), n = 11 (control). (E) Inflammatory fold induction in POWV LI9 whole brains were measured 15 dpi by qRT-PCR in comparison to control. For E: n = 3 (10, 20, 30, 40, 50 wk: 15 dpi), n = 3 (control).

We determined the minimal infectious dose required for LI9 lethality by s.c. infecting, or mock infecting, 50 wk old mice with 2 or 20 FFU of POWV LI9. Mice infected with 2 or 20 FFU of LI9, developed neurovirulent symptoms and 10-15% weight loss from 8-17 dpi (Supp. Fig. 1 C-E). Consistent with neuropathology, 50 wk old mice infected with either 2 or 20 FFU of LI9 succumbed to infection (10-16 dpi) with 50% or 90% mortality rates, respectively (Supp. Fig. 1D N=20/group). This data reveals that a minimal infectious dose of POWV LI9 (2-20 FFU) is highly lethal in aged mice.

Following peripheral inoculation of mice, POWV LI9 neuroinvasion was measured by qRT-PCR of viral RNA 5, 10 and 15 dpi (Fig. 1D). Despite age-dependent lethality, LI9 RNA was detected in murine brains from all age groups 5-15 dpi with the highest viral burdens at 10 dpi (10^4^-10^10^ copies/mL). By 15 dpi, viral RNA in the brain of 10, 20 and 30 wk mice decreased by >2 logs from 10 dpi levels (<10^4^ copies/mL). Brain RNA levels in 40 wk old mice decreased by 2-3 logs, but remained >10^5^ copies/mL. In contrast, in 50 wk old mouse brains, LI9 RNA levels were reduced by only ∼1 log from 10-15 dpi, but 10^8^ copies/mL of LI9 RNA was still present 15 dpi. This data suggests that POWV LI9 enters the CNS irrespective of murine age. However, in the brains of 10-30 wk old mice, LI9 is effectively cleared from the CNS, while 40-50 wk old mice either partially or completely fail to reduce viral CNS loads. These results suggest age-dependent differences in the viral load of LI9 infected brains 10-15 dpi that coincide with LI9 lethality in aged but not young mice.

Inflammation of the CNS contributes to neuronal demise, neuropathology and mortality^[13, 16, 17, 32]^. We assayed brain tissue of POWV LI9 infected 50 wk old mice for inflammatory chemokine and cytokine transcriptional responses 15 dpi (Fig. 1E). Consistent with increased viral load in the brain, pro-inflammatory chemokines IL-1β, CXCL10, CCL2 and CCL5 were highly induced in 50-wk-old mouse brains 15 dpi. These findings are consistent with POWV neuroinvasion, failure to clear virus from the CNS and the induction of neuroinflammatory responses that contribute to age-dependent neurovirulence.

### Neuropathology in POWV LI9 Infected Mice

To assess changes in brain pathology associated with lethal POWV LI9 neurovirulence, we evaluated formalin-fixed and H&E stained 50 wk old mouse brains 15 dpi for histological changes (Fig. 2). Analysis of POWV LI9 infected brains revealed severe neuropathology including: microgliosis, neuronal necrosis, spongiform encephalopathy, fibrosis and perivascular cuffing (Fig. 2A). Histologic lesions were present in several regions of the POWV LI9 infected brains including the pons, medulla, cerebellum, midbrain, cerebral cortex and brainstem (Fig. 2A). Scoring for spongiform encephalopathy (Figure 2B), microgliosis and neuronal necrosis (Supp. Fig. 2A-B) relative to controls, indicated the cerebellum and brainstem had the most severe pathology 15 dpi, followed by the pons and medulla.

**Figure 2.**
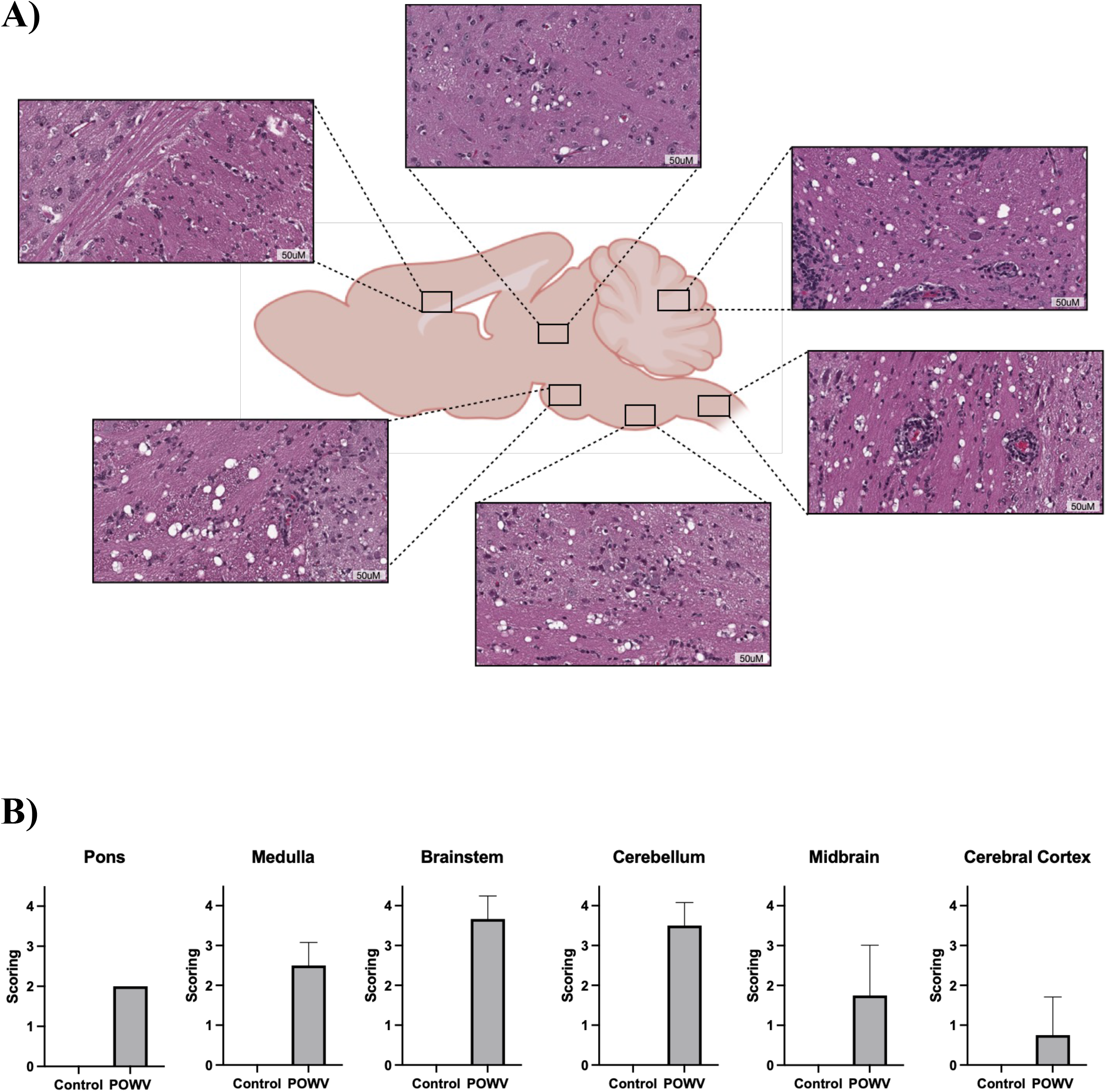
Distribution and severity of histologic lesions in POWV LI9 infected mice brains 15 dpi. (A-B) 50 wk old mice were inoculated with 2x10^3^ FFU of POWV LI9 or mock-infected with PBS by s.c. footpad injection. Sagittal cross-section of whole brains harvested at 15 dpi and H&E stained. (A) Representative images of POWV distribution throughout brain were taken. (B) H&E Brain sections were scored for spongiform encephalopathy severity on a scale of 0-4 compared to control. For A-B: n=4 (15 dpi), n=1 (control).

Kinetic analysis of 50 wk old LI9 infected brains revealed pathology by 5 dpi with clear histologic signs of inflammation, neuronal necrosis and spongiform encephalopathy (Fig. 3). A reduction in spongiform lesions was noted in LI9 infected brains between 5 and 10 dpi. Despite this, 10 dpi brains reveal inflammatory infiltrates and fibrosis that suggest microglial recruitment to sites of infection 5-10 dpi. By 15 dpi LI9 infected brains reveal new spongiform lesions, neuronal necrosis and persistent inflammatory infiltrates in brain parenchyma. Lesions developed from 10 to 15 dpi are consistent with sustained levels of POWV RNA and inflammatory transcripts from 10 to 15 dpi in 50 wk old brains (Fig. 1D-E). Analysis of a LI9 50 wk old mouse brain survivor (30 dpi), revealed prolonged damage and inflammation in the brain tissues (Supp. Fig. 3).

**Figure 3.**
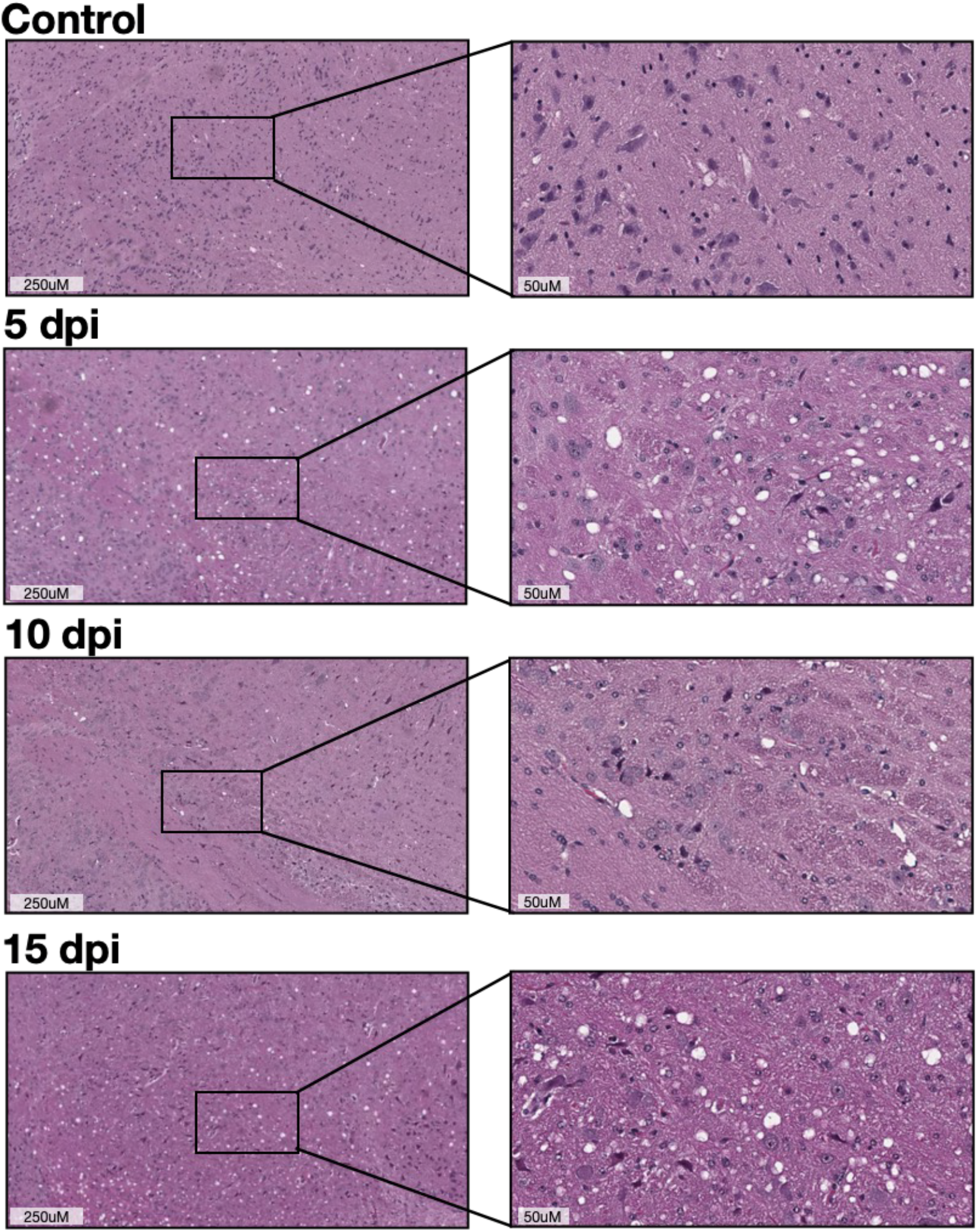
Hematoxylin and Eosin-Stained Sections of POWV LI9 Infected Mice Brains. 50 wk old mice were inoculated with 2x10^3^ FFU of POWV LI9 or mock-infected with PBS by s.c. footpad injection. Sagittal cross-section of whole brains harvested at 5, 10, 15 and 30 dpi and H&E stained. Representative images of Pons were taken. For 50 wk old: n=4 (5, 10, 15 dpi).

Neurons have been indicated as cellular targets of neurotropic FVs^[11, 12]^, and in TBEV infected neurons the presence of distinct punctate cytoplasmic structures and aberrant human neuronal dendrites were noted, without apoptosis^[9]^. Consistent with this, staining of LI9 infected 50 wk old brains with cresyl violet revealed neuronal Nissl bodies (5 dpi), associated with injury or disease. In contrast, alizarin red staining failed to detect intracranial calcium deposition in LI9 infected brains 5-15 dpi.

Neuronal damage and CNS pathology can be mediated directly by viral infection or indirectly by inflammatory responses within the CNS^[7]^. As an initial means of assessing immunopathology, we compared immunohistochemistry (IHC) of microglia and neurons in mock and POWV LI9 infected 50 wk old brains (5-15 dpi) (Fig. 4). Staining for microglia (Iba1) in POWV LI9 brains revealed microglial activation in the cerebral cortex, midbrain, cerebellum, pons, medulla and brainstem (Fig. 4A). By 5 dpi, activated microglia are present in POWV LI9 infected 50 wk old mice brain parenchyma and remain present 10 dpi (Fig. 4B). Consistent with prolonged CNS inflammation, IHC analysis revealed abundant activated microglia distributed throughout LI9 infected 50 wk old brains 15 dpi (Fig. 4A-B). Staining for neurons (NeuN) in 50 wk old POWV LI9 brains revealed a dramatic depletion of neurons within the medulla by 5 dpi (Fig. 4C), with neuronal loss coincident with medullary spongiform encephalopathy. H&E staining suggests the recovery of spongiform lesions within the brain by 10 dpi (Fig. 3), with NeuN positive neurons remaining at low levels 15 dpi (Fig. 4C). CNS findings are consistent with neuronal damage and prolonged inflammation, that fails to clear POWV from brains of 50 wk old mice, contributing to lethality.

**Figure 4.**
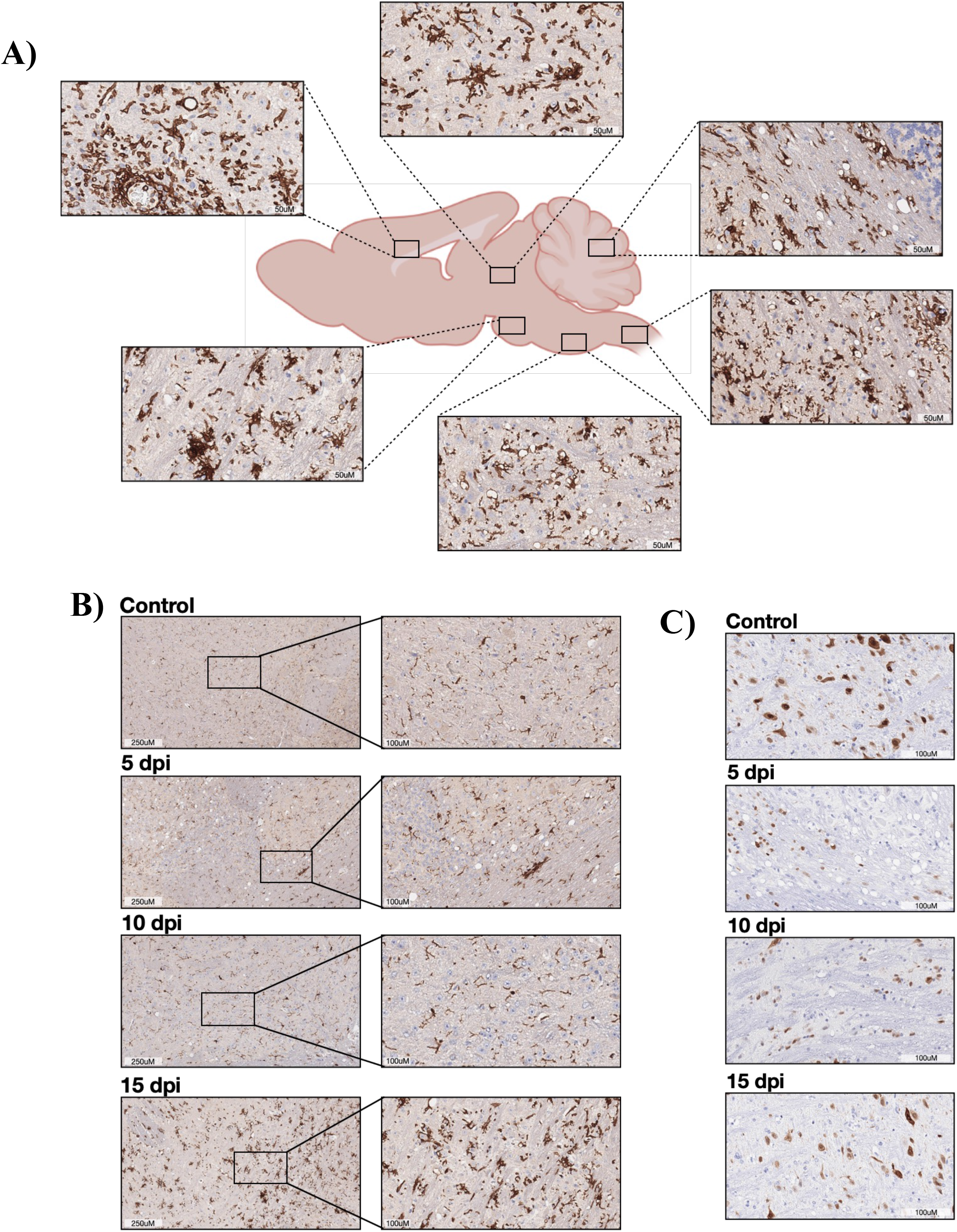
Iba1 or NeuN Stained Sections of POWV LI9 Infected Mice Brains. 50 wk old mice were inoculated with 2x10^3^ FFU of POWV LI9 or mock-infected with PBS by s.c. footpad injection. Sagittal cross-section of whole brains harvested at 5, 10, 15 and 30 dpi and immunohistochemical staining for Iba1 or NeuN was performed. Representative images were taken for (A) Global distribution of Iba1 staining 15 dpi throughout infected mouse brain. (B) Iba1 staining in Pons 5, 10 and 15 dpi. (C) NeuN staining in Medulla 5 dpi.

## Discussion

POWVs are rapidly transmitted in tick saliva, cause an acute febrile illness and, in severe cases, encephalitis with long term neurologic damage in patients^[1, 4, 33-42]^. POWV seroprevalence studies suggest that infections are largely undetected or undiagnosed in patients and this has limited human neuropathology studies. Improved clinical awareness and POWV testing, indicate that POWV incidence is increasing with the expanded geographic range of POWV vector species and shorter dormant seasons that exacerbate POWV spread^[43]^. There is little data on the mechanisms of POWV neuroinvasion and neurovirulence, or correlates of lethal POWV disease in humans, and currently there are no therapeutics or approved vaccines to prevent neurovirulent POWV disease.

Lineage II POWVs are thought to be more frequent causes of human disease, however, the pathology of fatal human encephalitis following POWV infections has been analyzed in only a few aged patients (age 63 to 79)^[44]^. POWV patient findings indicate profound cerebellar and brainstem involvement, with marked Purkinje cell loss, microgliosis, severe neuronal loss, T-cell infiltrates, viral RNA in brain and the lack of peripheral disease. Lethal human TBEV encephalitis is linked to aged individuals with no deaths in <40 yr olds and 1.5-3.5% lethality in 60-89 yr olds^[45, 46]^. Understanding POWV neurovirulence targets and responses is crucial to revealing age-dependent determinants of lethal encephalitis, and developing preventative and therapeutic approaches.

Murine models of POWV infection have been developed that establish lethal endpoints, analyze brain pathology and have been used for evaluating potentially protective POWV vaccines^[7, 27, 47]^. However, mortality rates in mice peripherally inoculated with POWV vary widely with a wide range of doses and suggested requirements for virulence. Prior studies were performed in 6-14 wk old mice with neuroadapted POWVs (LB/SP/IPS1) that were first serial passaged and selected for their ability to replicate in mouse brains^[27, 28, 48]^. POWV passage history and the age of mice used for experiments may partially explain varied lethality, dose independence, inocula required for neurovirulence (10^3^ to 10^6^), or suggested requirements for tick salivary gland extracts for spread to the CNS^[27, 28, 30]^.

To avoid neuroadaptive changes that may alter spread from peripheral sites to the CNS^[49, 50]^, we isolated POWV LI9 from deer ticks by inoculating VeroE6 cells with tick homogenates^[22]^. POWV LI9 is a Lin II POWV that is >99% identical to other Northeastern deer tick POWVs. *In vivo* POWV LI9 is neuroinvasive, causes clinical disease symptoms in C57BL/6 mice and elicits cross-reactive neutralizing POWV antibodies^[22]^. In initial experiments the fortuitous comparison of POWV LI9 infected 10 and 40 wk old mice suggested that lethality increased with murine age (Supp. Fig 1AB). Critical analysis of POWV in 10-50 wk old mice (N=10-20/group) revealed that murine lethality and weight loss were age-dependent. Aged, 50 wk old, mice had significantly higher fatality rates (80%) and lethality was reduced sequentially (40-10%) in 40>30>20>10 wk old mice and directly correlated with age. Minimal doses of POWV LI9 (2-20 FFUs) cause weight loss and 50-90% lethal disease in 50 wk old mice. These new novel findings reflect human POWV disease incidence in elderly patients^[2]^ and provide a murine model of age-dependent POWV LI9 neurovirulence. A novel neurovirulent tick-derived POWV, that directs age-dependent murine lethality and neuropathology, permits us to assess determinants of POWV neuroinvasion and neurovirulence at cell and systemic levels.

The distribution of POWV LI9 induced CNS pathology in 50 wk old mouse brains is similar to reports of human TBEV and POWV infections that cause CNS lesions throughout the brain^[7-9, 15, 16, 27, 30, 44, 51, 52]^. While we note some pathology in the midbrain and cerebral cortex, we find severe POWV LI9 induced lesions in the cerebellum and hindbrain region including the pons, medulla and brainstem. The distribution of POWV LI9 histopathological lesions in the brain demonstrate similarities of infection with human POWV pathology. However, in POWV LI9 infected mice, we detected POWV RNA in the brains of all mice, but in contrast neuroinvasion did not determine POWV lethality. Instead POWV lethality appears to be determined by differences in the ability of young vs aged mice to clear viral load within the brain^[7, 19, 53]^. In aged 40-50 wk old mouse brains POWV RNA was detected for a prolonged period, along with upregulated inflammatory transcripts and microglial infiltrates.

Histopathologic lesions in the CNS of LI9 infected aged mouse brains are consistent with severe CNS diseases and human POWV neuropathology findings. POWV infected 50 wk old mice demonstrate clear spongiform brain pathology that is associated with neurotropic infections and neurodegenerative brain disorders including Alzheimer’s and prion diseases^[54]^. Analysis of POWV infected aged brains revealed the early loss of neurons, microglial activation and inflammatory responses that are consistent with long term neurologic sequalae. In elderly individuals POWV neurovirulence is highly lethal but 50% of surviving patients also have long term severe neurological deficits consistent with spongiform neuropathology^[1, 2, 44]^.

Aging is associated with distinct changes in immune cell populations and function. Age-dependent changes in the CNS are linked to changes in the BBB and blood-cerebral-spinal-fluid-barrier (BCSFBs) that regulate immune cell infiltrates and direct neuroinflammatory T-cell responses that may be protective or contribute to neurologic damage^[19, 55-64]^. Old age also increases the risk of neurovirulent WNV illness 20-fold^[19]^ and in aged mice WNV increases viral burden in the CNS resulting from age-related defects in peripheral T-cells and cytokine signaling responses^[65, 66]^. Whether POWV neurovirulence is similarly immunoregulated in aged mice, remains to be determined but could reflect differences in the age-dependent clearance of POWV from brains.

POWV induced age-dependent CNS pathology may be a product of cellular damage activating inflammatory responses chemokine responses and cellular infiltrates^[7]^. Pro-inflammatory, IL-1β, CCL2, CCL5 and CXCL10 are upregulated by POWV infection *in vitro*^[22]^ and are observed *in vivo* in aged mouse brain responses to LI9 infection 15 dpi. IL-1β is a proinflamatory chemokine upregulated in neurodegenerative diseases, with pleotropic effects on apoptosis, CD8+ T cell recruitment and overall inflammation during encephalitic infections^[12, 17]^. Chemokines CCL2 and CCL5 positively or negatively modulate JEV neurovirulence^[67]^, and CXCL10 directs CD8+ T cell recruitment in WNV infection^[19]^. These responses could foster POWV clearance or exacerbate neuronal damage^[14, 19, 22, 64, 68-73]^ and responses of young versus aged brains may distinguish protective reactions from lethal outcomes.

The CNS response to FV infection includes activation of resident immune microglia, which play an essential role in viral clearance from the brain^[74, 75]^. Paradoxically, prolonged activation of inflammatory microglia is thought to induce or accelerate brain aging by interfering with repair and remodeling responses. Many neurotropic infections, cause low-grade neuroinflammation that is thought to contribute to brain aging and long-term neurologic sequalae^[76, 77]^. In aged individuals, microglia are thought to cause low-level state of inflammation compared to younger brains and in Alzheimer’s disease damage-associated microglia are adjacent to brain lesions and cause a severe proinflammatory state^[78]^. Consistent with prolonged inflammation, LI9 infected aged mouse brains express proinflammatory markers and demonstrate increased levels of activated microglia 5–15 dpi. This suggests that microglial antiviral responses may contribute to disease severity and long-term neurological sequalae observed in patients. Understanding microglial inflammatory responses, that fail to clear POWV from the CNS in aged mice, is critical for understanding human age-dependent POWV lethality.

Our findings provide an age-dependent lethal animal model to study POWV neuroinvasion and neurovirulence *in vivo*. We demonstrate that non-neuroadapted POWV LI9 infects spreads to the brain and is neuroinvasive following peripheral inoculation of both young and aged mice. Despite entry into young and old mouse brains, the lethality of LI9 is age-dependent and correlated with sustained viral load in the CNS of 50 wk old mice. Our results establish a role for age in lethal POWV disease that permit analysis of age-dependent changes which confer neurovirulence or protect young mice, and are likely to reveal mechanisms of severe human POWV disease in the elderly.

## MATERIALS AND METHODS

### Cells and Virus

VeroE6 cells (ATCC CRL 1586) were grown in DMEM (Dulbecco’s modified Eagle’s medium) supplemented with 5% heat-inactivated fetal bovine serum (FBS) and penicillin (100 mg/mL), streptomycin sulfate (100 mg/mL), and amphotericin B (50 mg/mL; Mediatech) at 37°C and 5% CO2. POWV strain LI9 (GenBank accession number: MZ576219) was isolated from infected *Ixodes scapularis* ticks via inoculation on Vero E6 cells and virus stocks were produced in Vero E6 cells^[22, 79]^. LI9 viral stocks (passage 3-4) were used to infect VeroE6 cells, isolate viral RNA and generate cDNA. All the work with infectious LI9 POWV was performed in a certified BSL3 facility at Stony Brook University.

### POWV LI9 Infection and Immunostaining

POWV LI9 was adsorbed to 60% confluent VeroE6 monolayers for 1 h. Following adsorption, monolayers were washed with PBS and grown in DMEM in 5% FBS. POWV LI9 titers were determined by serial dilution and infection of VeroE6 cells, quantifying infected cell foci at 24 hpi by immunoperoxidase staining with anti-POWV hyperimmune mouse ascites fluid (HMAF; 1:5,000 [ATCC]), horseradish peroxidase (HRP)-labeled anti-mouse IgG (1:2,000; KPL-074-1806), and 3-amino-9-ethylcarbazole (AEC) staining (60).

### Murine inoculation

C57BL/6J mice (10-50 weeks old) were purchased from Jackson Laboratory. Mice were anesthetized via intraperitoneal injection with 100 mg·ml^-1^ of ketamine and 20 mg·ml^-1^ of xylazine per kilogram of body weight. Animals were infected via subcutaneous footpad injection with up to 2 × 10^3^ FFU POWV or buffer only control in a volume of 20 μl. Mice were monitored daily for signs of disease and daily weight loss. At 5, 10, 15, 30 dpi mice were euthanized by CO_2_ inhalation for sera and brain collection.

### Biosafety and Biosecurity

Animal research was performed in accordance with institutional guidelines following experimental protocol review, approval, and supervision by the Institutional Biosafety Committee and the Institutional Animal Care and Use Committee at Stony Brook University. Animals were managed by the Division of Laboratory Animal Resources (Stony Brook University), which is accredited by the American Association for Accreditation of Laboratory Animal Care and the Department of Health and Human Services. Animals were maintained in accordance with the applicable portions of the Animal Welfare Act and the DHHS ‘Guide for the Care and Use of Laboratory Animals.’ Veterinary care was under the direction of full-time resident veterinarians boarded by the American College of Laboratory Animal Medicine. Experiments with infectious POWV were performed in biosafety level 3 containment (SBU ABSL3 facility).

### Histopathological Analysis

Brains were harvested postmortem, fixed in neutral buffered formalin for 7 days, dehydrated with 70% ethanol for 24 h and paraffin embedded. Formalin-fixed paraffin-embedded (FFPE) brain tissues were routinely sectioned (10 μm thickness) and stained by the Stony Brook Research Histology Core Lab or by HistoWiz, Inc. Histochemical staining with hematoxylin and eosin (H&E) was performed used for observation of cell nuclei, cytoplasm, and tissue architecture. Immunohistochemical staining with anti-Iba1 antibody (HistoWiz Catalog # Wako 019-19741) was used for identification and analysis of activated microglia. Immunohistochemical staining with anti-NeuN antibody (HistoWiz, Catalog # ab177487) was used for identification and analysis of neurons. Immunohistochemical staining with anti-POWV HMAF (ATCC V-518-711-562) was used for identification and analysis of POWV infected cells.

Stained tissue sections were dehydrated by immersing samples in a series of ethanol solutions of increasing concentrations to xylene and mounted using a nonaqueous mounting medium, after which slides were digitized by Stony Brook Research Histology Core Lab or HistoWiz, Inc. Blind, and analyzed using the latest version of QuPath software (https://qupath.github.io).

### RNA Extraction and qRT-PCR analyses

Mock or POWV-infected mice were euthanized as described above and brains were harvested in TRIzol LS Reagent (Invitrogen, Life Technologies) according to manufacturer’s recommendations. Tissues were homogenized and chloroform was added to the tissue homogenates at a volume recommended by the TRIzol reagent protocol. The samples were mixed vigorously for 15 seconds, incubated at room temperature for 3 minutes, and centrifuged at 12,000 x g for 15 minutes at 4°C. The upper aqueous phase was collected, and one volume of 70% ethanol was added to each sample and mixed (thang). RNA was extracted from samples using Monarch RNA Cleanup Kit (NEB T2030L) and quantified using a Nanodrop 2000 Spectrophotometer 2000.

To define viral loads and inflammatory transcripts quantitative real-time PCR was performed on RNAs from purified brain homogenates from mock or POWV infected mice. cDNA synthesis was performed using a Transcriptor first-strand cDNA synthesis kit (Roche) using random hexamers as primers (25°C for 10 min, 50°C for 60 min, and 90°C for 5 min). Quantification of viral loads in brain tissue were determined using paired NS5 specific LI9 primers (Forward-GAAACAATACTCAGAATCATG, Reverse-AAGCCGCTGATCCAGTGGCA). Viral burden is expressed on a Log scale as viral RNA copies per milliliter after comparison with a standard curve produced using serial 10-fold dilutions of POWV RNA. qRT-PCR transcript primers for IL-1Beta (Forward-GGAGAACCAAGCAACGACAAAATA, Reverse-TGGGGAACTCTGCAGACTCAAAC), CCL2 (Forward-TCAGCCAGATGCAGTTAACG, Reverse-CTCTCTTGAGCTTGGTGACA), CCL5 (Forward-CAAGTGCTCCAATCTTGCAG, Reverse-CCTCTATCCTAGCTCATCTCCA), CXCL10 (Forward-AGTGCTGCCGTCATTTTCTG, Reverse-ATTCTCACTGGCCCGTCAT) were designed according to the NCBI gene database, with 60°C annealing profiles. Genes were analyzed using PerfeCTa SYBR green SuperMix (Quanta Biosciences) on a Bio-Rad C1000 Touch system with a CFX96 optical module (Bio-Rad). Responses were normalized to internal glyceraldehyde-3-phosphate dehydrogenase (GAPDH) mRNA levels, and the fold induction was calculated using the threshold cycle (2^ddCT^) method for differences between mock- and POWV-infected RNA levels at each time point.

### Neutralizing Antibody Assays

Neutralizing antibodies present in POWV LI9, or mock-infected mouse sera 30 dpi were determined by serial tenfold dilution and addition to ∼500 FFU of POWV LI9. Virus was adsorbed to VeroE6 cells and POWV infected cells were quantitated 36 hpi by immunoperoxidase staining as above, and reciprocal dilutions conferring a 50% reduction in infected cells versus controls was determined.

## Statistical analysis

All of the details regarding the statistical analysis of the data can be found in the figure legends. The significance of the results described in this study was determined by use of a number of different statistical analyses performed with Prism 6 software (GraphPad Software, Inc.; https://www.graphpad.com). P values of less than 0.05 were considered statistically significant.

## ACKNOWLEDGEMENTS

We thank Jorge Benach, Dan Salamongo, Nicholas Carpino, and Jeronimo Cello for manuscript feedback and supportive discussions on tick-borne diseases and neuropathogenesis, and Stephen Gaudino, Smruti Mishra and Luke Helminiak for guidance on immunohistochemistry and animal monitoring and sample retrieval. This work was supported by funding from a DOD TBDRP Idea Development Award W81XWH2210702, National Institutes of Health grants: NIAID R01AI12901005, R21AI13173902, R21AI15237201, RO1AI027044, T32AI007539, an IRACDA Post-Doctoral Award and a Stony Brook University Seed Grant. The funders had no role in study design, data collection and interpretation or the decision to submit the work for publication. We declare no conflict of interest.

## FIGURE LEGENDS

**Supplemental Figure 1. Aged Mice are Susceptible to POWV LI9. (**A-B) 10-, and 40-wk-old mice were inoculated with 10^3^ FFU of POWV LI9 by subcutaneous (s.c.) footpad injection. (A) Mice were weighed daily, and weights were expressed as a percentage of body weight prior to infection. Results are the mean +/- standard error of the mean (SEM) of the indicated number of mice per group and were analyzed by use of a log rank test. **, *P <* 0.01; *, *P <* 0.05. (B) Lethality was monitored for 15 days. For (A-B): n = 4 (control), n = 4 (10wk), n = 4 (40wk). (C-E) 50 wk old mice were inoculated with 2 or 20 FFU of POWV LI9 or mock-infected with PBS by subcutaneous (s.c.) footpad injection. (C) Mice were weighed daily, and weights were expressed as a percentage of body weight prior to infection. Results are the mean +/- standard error of the mean (SEM) of the indicated number of mice per group and were analyzed by use of a log rank test ****, *P <* 0.0001; **, *P <* 0.01; *, *P <* 0.05. (D) Lethality was monitored for 30 days. For (C-D): n = 20 (2 FFU), n = 20 (20 FFU), n = 20 (mock-infected control). (E) Mouse sera was collected 30 dpi and analyzed for neutralizing antibody responses by Western blot detection of POWV LI9 infected VeroE6 cell lysates (3 dpi). For (E): n = 3 (2 FFU), n = 1 (20 FFU), n = 3 (200 FFU), n = 3 (2000 FFU).

**Supplemental Figure 2. Distribution and severity of neuronal necrosis, and microgliosis in POWV LI9 infected mice brains 15 dpi**. (A-B) 50 wk old mice were inoculated with 2x10^3^ FFU of POWV LI9 or mock-infected with PBS by s.c. footpad injection. Sagittal cross-section of whole brains harvested at 15 dpi and H&E stained. H&E sections were scored for (A) neuronal necrosis or (B) microgliosis severity on a scale of 0-4 compared to control. For (A-B): n=4 (15 dpi), n=1 (control).

**Supplemental Figure 3. Hematoxylin and Eosin-Stained Sections of POWV LI9 Infected 50 wk old Suvivor Mouse Brain**. 50 wk old mice were inoculated with 2x10^3^ FFU of POWV LI9 or mock-infected with PBS by s.c. footpad injection. Sagittal cross-section of whole brains harvested at 30 dpi and H&E stained. Representative images of Pons were taken. For 50 wk old: n=1 (30 dpi, 3 additional mice did not survive).

**Supplemental Figure 4. POWV Antigen-Stained Sections of Intracranial Infected POWV LI9 Pup Brain**. C57BL/6 pups were inoculated with 2x10^2^ FFU of POWV LI9 or mock-infected with PBS by s.c. footpad injection. Sagittal cross-section of whole brains harvested at 3 dpi and immunhistochemical staining with POWV HMAF (ATCC) was performed. Representative images of cerebral cortex were taken.

## REFERENCES

1. Kemenesi, G. and K. Banyai, Tick-Borne Flaviviruses, with a Focus on Powassan Virus. Clin Microbiol Rev, 2019. 32(1).

2. Campbell, O. and P.J. Krause, The emergence of human Powassan virus infection in North America. Ticks Tick Borne Dis, 2020. 11(6): p. 101540.

3. Bonet, R., I. Vakonakis, and I.D. Campbell, Characterization of 14-3-3-zeta Interactions with integrin tails. J Mol Biol, 2013. 425(17): p. 3060–72.

4. Ebel, G.D. and L.D. Kramer, Short report: duration of tick attachment required for transmission of powassan virus by deer ticks. Am J Trop Med Hyg, 2004. 71(3): p. 268–71.

5. Ebel, G.D., et al., A focus of deer tick virus transmission in the northcentral United States. Emerg Infect Dis, 1999. 5(4): p. 570–4.

6. McLean, D.M., S.R. Ladyman, and K.W. Purvin-Good, Westward extension of Powassan virus prevalence. Can Med Assoc J, 1968. 98(20): p. 946–9.

7. Hayasaka, D., et al., Mortality following peripheral infection with tick-borne encephalitis virus results from a combination of central nervous system pathology, systemic inflammatory and stress responses. Virology, 2009. 390(1): p. 139–50.

8. Nagata, N., et al., The pathogenesis of 3 neurotropic flaviviruses in a mouse model depends on the route of neuroinvasion after viremia. J Neuropathol Exp Neurol, 2015. 74(3): p. 250–60.

9. Nelson, J., et al., Powassan Virus Induces Structural Changes in Human Neuronal Cells In Vitro and Murine Neurons In Vivo. Pathogens, 2022. 11(10).

10. Abbink, P., et al., Protective efficacy of multiple vaccine platforms against Zika virus challenge in rhesus monkeys. Science. 353(6304): p. 1129–32.

11. Chambers, T.J. and M.S. Diamond, Pathogenesis of flavivirus encephalitis. Adv Virus Res, 2003. 60: p. 273–342.

12. Fares, M., et al., Pathological modeling of TBEV infection reveals differential innate immune responses in human neurons and astrocytes that correlate with their susceptibility to infection. J Neuroinflammation, 2020. 17(1): p. 76.

13. Diamond, M.S., et al., The host immunologic response to West Nile encephalitis virus. Front Biosci (Landmark Ed), 2009. 14: p. 3024–34.

14. Durrant, D.M., et al., CCR5 limits cortical viral loads during West Nile virus infection of the central nervous system. J Neuroinflammation. 12: p. 233.

15. Gelpi, E., et al., Visualization of Central European tick-borne encephalitis infection in fatal human cases. J Neuropathol Exp Neurol, 2005. 64(6): p. 506–12.

16. Gelpi, E., et al., Inflammatory response in human tick-borne encephalitis: analysis of postmortem brain tissue. J Neurovirol, 2006. 12(4): p. 322–7.

17. Ruzek, D., et al., CD8+ T-cells mediate immunopathology in tick-borne encephalitis. Virology, 2009. 384(1): p. 1–6.

18. Krow-Lucal, E.R., et al., Powassan Virus Disease in the United States, 2006-2016. Vector Borne Zoonotic Dis, 2018. 18(6): p. 286–290.

19. Klein, R.S., et al., Neuronal CXCL10 directs CD8+ T-cell recruitment and control of West Nile virus encephalitis. J Virol, 2005. 79(17): p. 11457–66.

20. Krawczuk, K., et al., Comparison of tick-borne encephalitis between children and adults-analysis of 669 patients. J Neurovirol, 2020. 26(4): p. 565–571.

21. Beasley, D.W., et al., Nucleotide sequencing and serological evidence that the recently recognized deer tick virus is a genotype of Powassan virus. Virus Res, 2001. 79(1-2): p. 81–9.

22. Conde, J.N., et al., Powassan Viruses Spread Cell to Cell during Direct Isolation from Ixodes Ticks and Persistently Infect Human Brain Endothelial Cells and Pericytes. J Virol, 2022. 96(1): p. e0168221.

23. Pesko, K.N., et al., Molecular epidemiology of Powassan virus in North America. J Gen Virol, 2010. 91(Pt 11): p. 2698–705.

24. Anderson, J.F. and P.M. Armstrong, Prevalence and genetic characterization of Powassan virus strains infecting Ixodes scapularis in Connecticut. Am J Trop Med Hyg, 2012. 87(4): p. 754–9.

25. VanBlargan, L.A., et al., Broadly neutralizing monoclonal antibodies protect against multiple tick-borne flaviviruses. J Exp Med, 2021. 218(5).

26. VanBlargan, L.A., et al., An mRNA Vaccine Protects Mice against Multiple Tick-Transmitted Flavivirus Infections. Cell Rep, 2018. 25(12): p. 3382–3392 e3.

27. Hermance, M.E., et al., Development of a small animal model for deer tick virus pathogenesis mimicking human clinical outcome. PLoS Negl Trop Dis, 2020. 14(6): p. e0008359.

28. Hermance, M.E. and S. Thangamani, Tick Saliva Enhances Powassan Virus Transmission to the Host, Influencing Its Dissemination and the Course of Disease. J Virol, 2015. 89(15): p. 7852–60.

29. Holbrook, M.R., et al., An animal model for the tickborne flavivirus--Omsk hemorrhagic fever virus. J Infect Dis, 2005. 191(1): p. 100–8.

30. Santos, R.I., et al., Salivary gland extract from the deer tick, Ixodes scapularis, facilitates neuroinvasion by Powassan virus in BALB/c mice. Sci Rep, 2021. 11(1): p. 20873.

31. Mlera, L., et al., Modeling Powassan virus infection in Peromyscus leucopus, a natural host. PLoS Negl Trop Dis, 2017. 11(1): p. e0005346.

32. Pierson, T.C. and M.S. Diamond, The continued threat of emerging flaviviruses. Nat Microbiol, 2020. 5(6): p. 796–812.

33. McLean, D. and W.L. Donohue, Powassan virus: isolation of virus from a fatal case of encephalitis. Can Med Assoc J, 1959. 80(9): p. 708–11.

34. Sanchez-Vicente, S., et al., Polymicrobial Nature of Tick-Borne Diseases. mBio, 2019. 10(5).

35. Fatmi, S.S., R. Zehra, and D.O. Carpenter, Powassan Virus-A New Reemerging Tick-Borne Disease. Front Public Health, 2017. 5: p. 342.

36. Rossier, E., R.J. Harrison, and B. Lemieux, A case of Powassan virus encephalitis. Can Med Assoc J, 1974. 110(10): p. 1173–4 passim.

37. Feder, H.M., et al., Powassan Virus Encephalitis Following Brief Attachment of Connecticut Deer Ticks. Clin Infect Dis, 2020.

38. Campagnolo, E.R., et al., Evidence of Powassan/deer tick virus in adult black-legged ticks (Ixodes scapularis) recovered from hunter-harvested white-tailed deer (Odocoileus virginianus) in Pennsylvania: A public health perspective. Zoonoses Public Health, 2018. 65(5): p. 589–594.

39. Hermance, M.E. and S. Thangamani, Powassan Virus: An Emerging Arbovirus of Public Health Concern in North America. Vector Borne Zoonotic Dis, 2017. 17(7): p. 453–462.

40. Khan, A.M., et al., Powassan virus encephalitis, severe babesiosis and lyme carditis in a single patient. BMJ Case Rep, 2019. 12(11).

41. Robich, R.M., et al., Prevalence and Genetic Characterization of Deer Tick Virus (Powassan Virus, Lineage II) in Ixodes scapularis Ticks Collected in Maine. Am J Trop Med Hyg, 2019. 101(2): p. 467–471.

42. Mihara, M., A Histopathologic Study of the Human Skin in the Early Stage After a Tick Bite: A Special Reference to Cutaneous Tissue Reaction to the Cement Substance of Tick Saliva. Yonago Acta Med, 2017. 60(3): p. 186–199.

43. Bouchard, C., et al., N Increased risk of tick-borne diseases with climate and environmental changes. Can Commun Dis Rep, 2019. 45(4): p. 83–89.

44. Normandin, E., et al., Powassan Virus Neuropathology and Genomic Diversity in Patients With Fatal Encephalitis. Open Forum Infect Dis, 2020. 7(10): p. ofaa392.

45. Varnaite, R., S. Gredmark-Russ, and J. Klingstrom, Deaths from Tick-Borne Encephalitis, Sweden. Emerging Infectious Diseases, 2022. 28(7): p. 1471–1474.

46. Bogovic, P. and F. Strle, Tick-borne encephalitis: A review of epidemiology, clinical characteristics, and management. World Journal of Clinical Cases, 2015. 3(5): p. 430–441.

47. King, N.J., et al., Immunopathology of flavivirus infections. Immunol Cell Biol, 2007. 85(1): p. 33–42.

48. Santos, R.I., et al., Spinal Cord Ventral Horns and Lymphoid Organ Involvement in Powassan Virus Infection in a Mouse Model. Viruses, 2016. 8(8).

49. Dubuisson, J., et al., Genetic determinants of Sindbis virus neuroinvasiveness. J Virol, 1997. 71(4): p. 2636–46.

50. Tucker, P.C., et al., Viral determinants of age-dependent virulence of Sindbis virus for mice. J Virol, 1993. 67(8): p. 4605–10.

51. Takashima, I., et al., A case of tick-borne encephalitis in Japan and isolation of the the virus. J Clin Microbiol, 1997. 35(8): p. 1943–7.

52. Dumpis, U., D. Crook, and J. Oksi, Tick-borne encephalitis. Clinical Infectious Diseases, 1999. 28(4): p. 882–890.

53. Palus, M., et al., Mice with different susceptibility to tick-borne encephalitis virus infection show selective neutralizing antibody response and inflammatory reaction in the central nervous system. J Neuroinflammation, 2013. 10: p. 77.

54. Broxmeyer, L., Thinking the unthinkable: Alzheimer’s, Creutzfeldt-Jakob and Mad Cow disease: the age-related reemergence of virulent, foodborne, bovine tuberculosis or losing your mind for the sake of a shake or burger. Med Hypotheses, 2005. 64(4): p. 699–705.

55. Alisch, J.S.R., et al., Characterization of Age-Related Differences in the Human Choroid Plexus Volume, Microstructural Integrity, and Blood Perfusion Using Multiparameter Magnetic Resonance Imaging. Front Aging Neurosci, 2021. 13: p. 734992.

56. Baruch, K., et al., Aging. Aging-induced type I interferon response at the choroid plexus negatively affects brain function. Science, 2014. 346(6205): p. 89–93.

57. Dani, N., et al., A cellular and spatial map of the choroid plexus across brain ventricles and ages. Cell, 2021. 184(11): p. 3056–3074 e21.

58. Llovera, G., et al., The choroid plexus is a key cerebral invasion route for T cells after stroke. Acta Neuropathol, 2017. 134(6): p. 851–868.

59. Saunders, N.R., et al., The choroid plexus: a missing link in our understanding of brain development and function. Physiol Rev, 2023. 103(1): p. 919–956.

60. Schwerk, C., et al., The choroid plexus-a multi-role player during infectious diseases of the CNS. Front Cell Neurosci, 2015. 9: p. 80.

61. Tahira, A., et al., Are the 50’s, the transition decade, in choroid plexus aging? Geroscience, 2021. 43(1): p. 225–237.

62. Thompson, D., C.A. Brissette, and J.A. Watt, The choroid plexus and its role in the pathogenesis of neurological infections. Fluids Barriers CNS, 2022. 19(1): p. 75.

63. Lowerison, M.R., et al., Aging-related cerebral microvascular changes visualized using ultrasound localization microscopy in the living mouse. Sci Rep, 2022. 12(1): p. 619.

64. Funk, K.E., et al., Decreased antiviral immune response within the central nervous system of aged mice is associated with increased lethality of West Nile virus encephalitis. Aging Cell, 2021. 20(8): p. e13412.

65. Richner, J.M., et al., Age-Dependent Cell Trafficking Defects in Draining Lymph Nodes Impair Adaptive Immunity and Control of West Nile Virus Infection. PLoS Pathog, 2015. 11(7): p. e1005027.

66. Brien, J.D., et al., Key role of T cell defects in age-related vulnerability to West Nile virus. J Exp Med, 2009. 206(12): p. 2735–45.

67. Kim, J.H., et al., CCL2, but not its receptor, is essential to restrict immune privileged central nervous system-invasion of Japanese encephalitis virus via regulating accumulation of CD11b(+) Ly-6C(hi) monocytes. Immunology, 2016. 149(2): p. 186–203.

68. Vidana, B., et al., West Nile Virus spread and differential chemokine response in the central nervous system of mice: Role in pathogenic mechanisms of encephalitis. Transbound Emerg Dis, 2020. 67(2): p. 799–810.

69. Stefanik, M., et al., Characterisation of Zika virus infection in primary human astrocytes. BMC Neurosci, 2018. 19(1): p. 5.

70. Zhang, X., et al., Tick-borne encephalitis virus induces chemokine RANTES expression via activation of IRF-3 pathway. J Neuroinflammation, 2016. 13(1): p. 209.

71. Zhang, F., et al., PD1(+)CCR2(+)CD8(+) T Cells Infiltrate the Central Nervous System during Acute Japanese Encephalitis Virus Infection. Virol Sin, 2019. 34(5): p. 538–548.

72. Berger, M.M., et al., Hypoxia impairs systemic endothelial function in individuals prone to high-altitude pulmonary edema. Am J Respir Crit Care Med, 2005. 172(6): p. 763–7.

73. Mladinich, M.C., et al., Blockade of Autocrine CCL5 Responses Inhibits Zika Virus Persistence and Spread in Human Brain Microvascular Endothelial Cells. mBio, 2021: p. e0196221.

74. Hatton, C.F. and C.J.A. Duncan, Microglia Are Essential to Protective Antiviral Immunity: Lessons From Mouse Models of Viral Encephalitis. Front Immunol, 2019. 10: p. 2656.

75. Ransohoff, R.M. and A.E. Cardona, The myeloid cells of the central nervous system parenchyma. Nature, 2010. 468(7321): p. 253–62.

76. Maximova, O.A. and A.G. Pletnev, Flaviviruses and the Central Nervous System: Revisiting Neuropathological Concepts. Annu Rev Virol, 2018. 5(1): p. 255–272.

77. Maximova, O.A., et al., Comparative neuropathogenesis and neurovirulence of attenuated flaviviruses in nonhuman primates. J Virol, 2008. 82(11): p. 5255–68.

78. Edler, M.K., I. Mhatre-Winters, and J.R. Richardson, Microglia in Aging and Alzheimer’s Disease: A Comparative Species Review. Cells, 2021. 10(5).

79. Conde, J.N., et al., NS5 Sumoylation Directs Nuclear Responses That Permit Zika Virus To Persistently Infect Human Brain Microvascular Endothelial Cells. J Virol, 2020. 94(19).

