## Supplementary figures and images for "Age-dependent Powassan Virus Lethality and Neuropathogenesis in Mice"

### Supplemental Fig 1-4

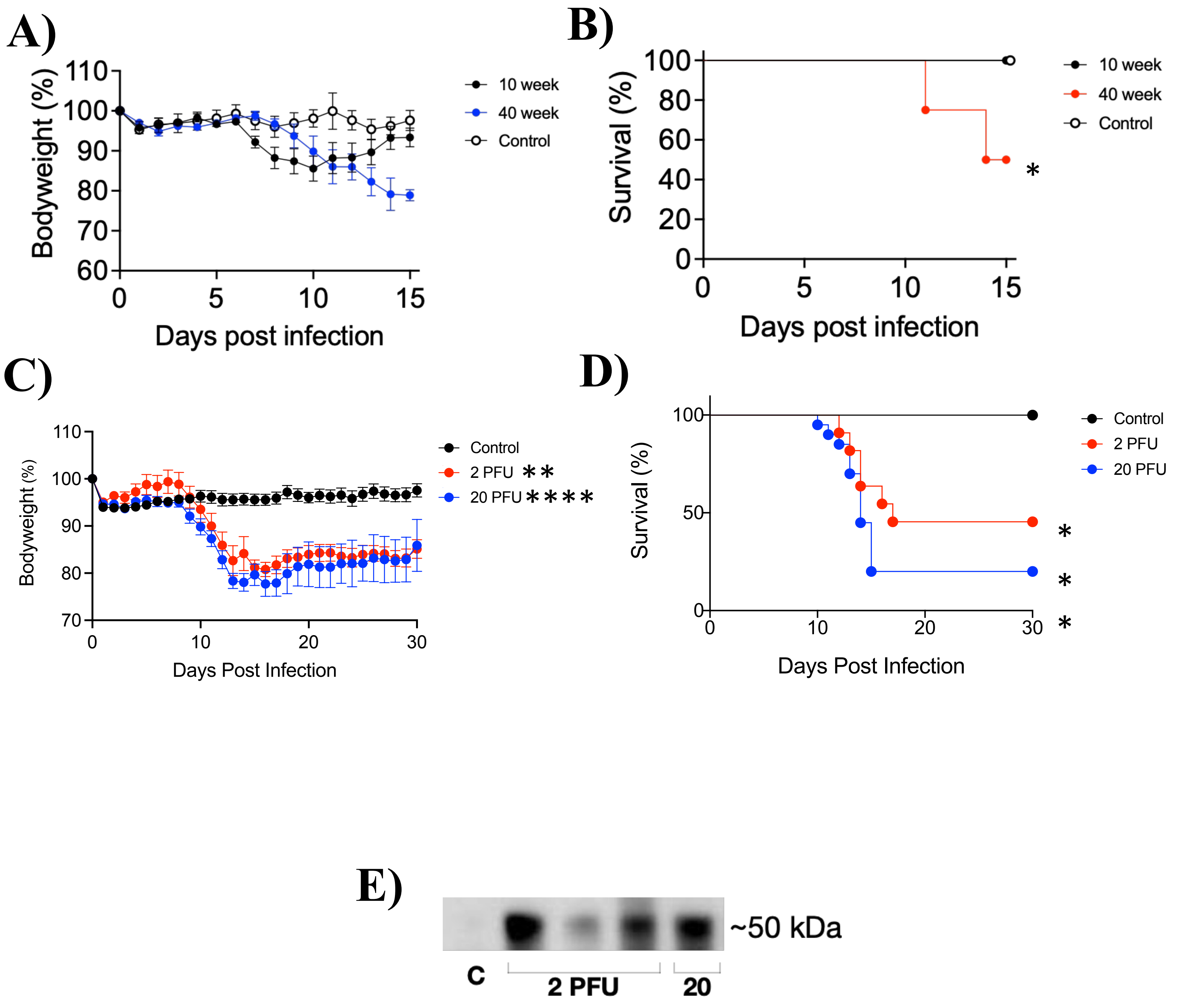


#### Supplemental Figure 1

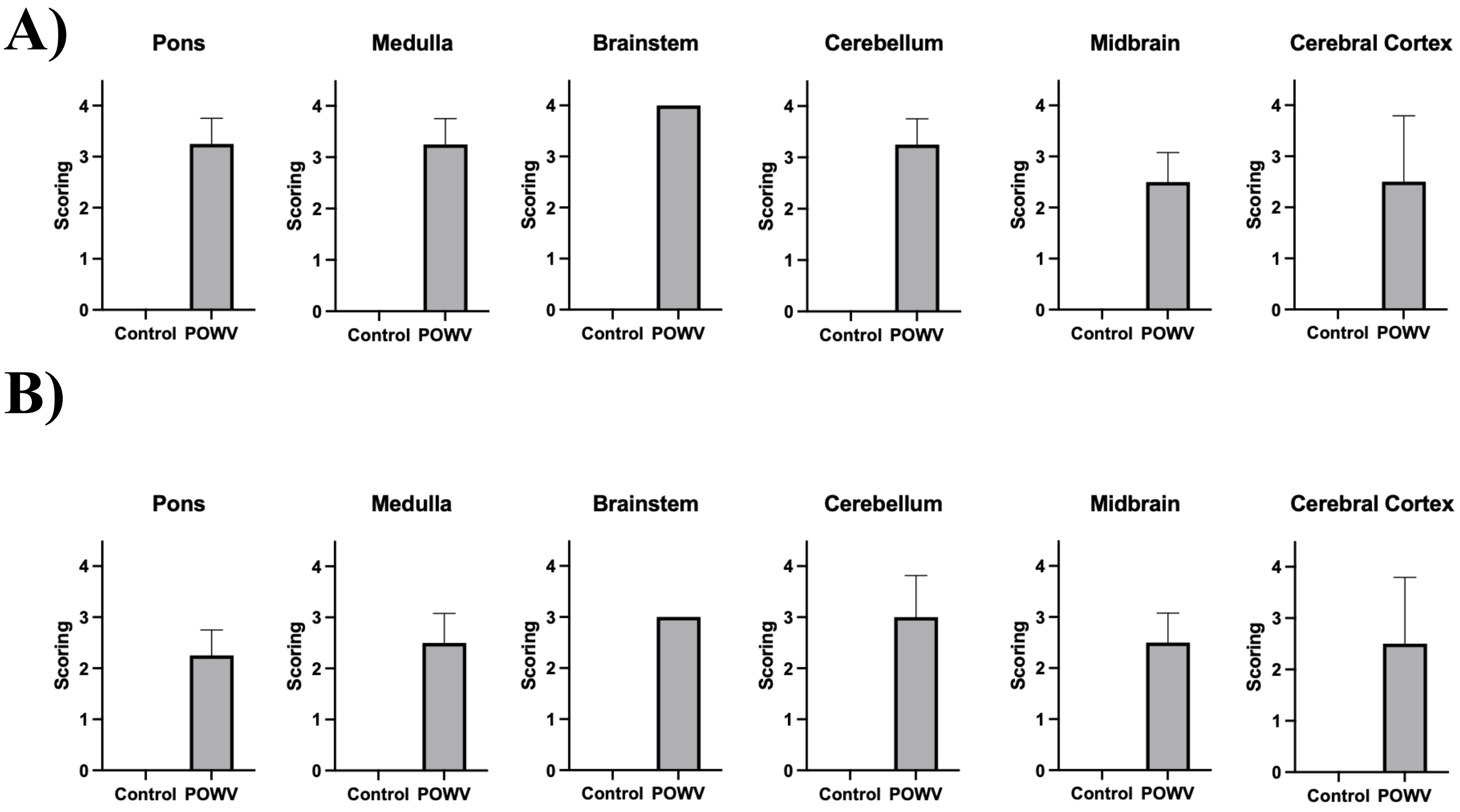


**Supplemental Figure 2**

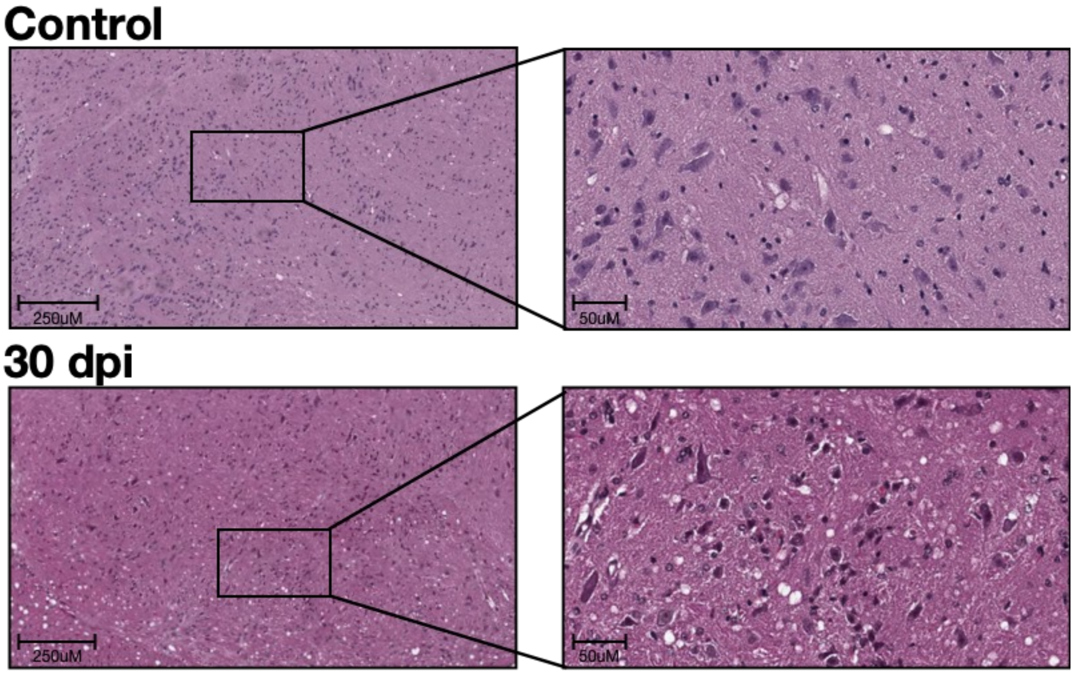


#### Supplemental Figure 3

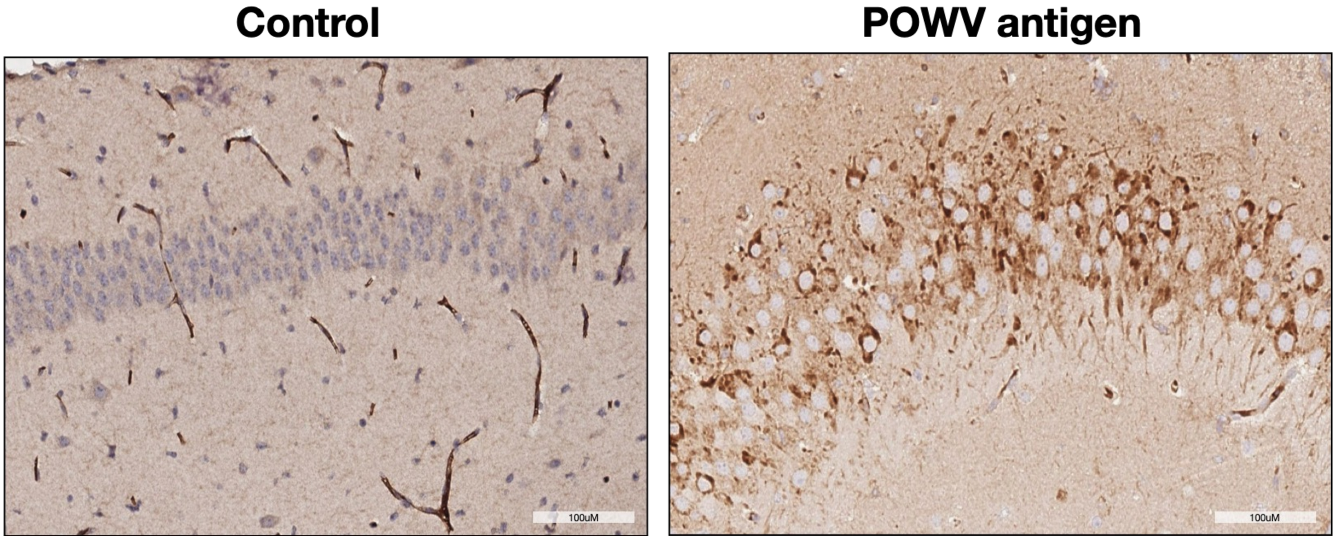


Supplemental Figure 4
